# *Bacteroides thetaiotaomicron* outer membrane vesicles modulate virulence of *Shigella flexneri*

**DOI:** 10.1101/2022.08.24.505214

**Authors:** Nicholas L. Xerri, Shelley M. Payne

## Abstract

The role of the gut microbiota in the pathogenesis of *Shigella flexneri* remains largely unknown. To understand the impact of the gut microbiota on *S. flexneri* virulence, we examined the effect of interspecies interactions with *Bacteroides thetaiotaomicron* (*Bt*), a prominent member of the gut microbiota, on *S. flexneri* invasion. When grown in *Bt* conditioned medium, *S. flexneri* showed reduced invasion of human epithelial cells. This decrease in invasiveness of *S. flexneri* resulted from a reduction in the level of *S. flexneri*’s master virulence regulator VirF. Reduction of VirF corresponded with a decrease in expression of a secondary virulence regulator *virB*, as well as expression of *S. flexneri* virulence genes required for invasion, intracellular motility, and spread. Repression of *S. flexneri* virulence factors by *Bt* conditioned medium was not caused by either a secreted metabolite or protein, but rather, was due to the presence of *Bt* outer membrane vesicles (OMVs) in the conditioned medium. The addition of purified *Bt* OMVs to *S. flexneri* growth medium recapitulated the inhibitory effects of *Bt* conditioned medium on invasion, virulence gene expression, and virulence protein levels. Total lipids extracted from either *Bt* cells or *Bt* OMVs also recapitulated the effects of *Bt* condition medium on expression of the *S. flexneri* virulence factor IpaC, indicating that *Bt* OMV lipids, rather than a cargo contained in the vesicles, are the active factor responsible for the inhibition of *S. flexneri* virulence.

**Importance:** *Shigella flexneri* is the causative agent of bacillary dysentery in humans. *Shigella* spp. are one of the leading causes of diarrheal morbidity and mortality, especially among children in low and middle-income countries. The rise of antimicrobial resistance combined with the lack of an effective vaccine for *Shigella* heightens the importance of studies aimed at better understanding previously uncharacterized aspects of *Shigella* pathogenesis. Here, we show that conditioned growth medium from the commensal bacteria *Bacteroides thetaiotaomicron* represses the invasion of *S. flexneri*. This repression is due to the presence of *B. thetaiotaomicron* outer membrane vesicles. These findings establish a role for interspecies interactions with a prominent member of the gut microbiota in modulating the virulence of *S. flexneri* and identify a novel function of outer membrane vesicles in interbacterial signaling between members of the gut microbiota and an enteric pathogen.

## Introduction

*Shigella flexneri* is an enteric pathogen that causes bacillary dysentery in humans (1). After being ingested, *S. flexneri* passes through the stomach and small intestine before entering the colon, where the expression of invasion genes (2) encoded on its virulence plasmid (3) enables it to invade the colonic epithelium (4). Coordination of invasion, vacuole lysis, and intercellular spread by *S. flexneri* relies on the timed expression of a complex array of virulence genes (5). The genes involved in this virulence process are controlled by a master transcriptional regulator VirF, which directly regulates *icsA*, a virulence gene encoding a protein necessary for *S. flexneri* motility within host cells, and *virB*, a gene encoding a secondary transcriptional regulator required for virulence (6, 7). VirB, in turn, regulates a variety of type three secretion system (T3SS) and invasion genes such as *ipaA, B, C*, and *D* (8).

Prior to gaining access to the intestinal epithelium for invasion, *S. flexneri* encounters a variety of potential obstacles in the colon, including the trillions of established bacteria that make up the human gut microbiota (9). However, the role of the gut microbiota in *S. flexneri* pathogenesis remains largely unknown. While conventional guinea pigs and mice are resistant to colonization by *S. flexneri*, germ-free animals are susceptible to colonization, indicating a possible role for the microbiota in resistance to infection (10). Monoassociating germ-free animals with *Escherichia coli* prior to challenge with *S. flexneri* restores resistance to *S. flexneri* colonization, but monoassociating mice with *Bacteroides*, a prominent member of the human gut microbiota, has no effect on *S. flexneri* colonization. Interestingly, di-associating mice with both *E. coli* and *Bacteroides* has the largest impact on *S. flexneri* colonization, causing *S. flexneri* to reduce to undetectable levels in the colon (10–12). Together, these studies suggest that the gut microbiota may function in preventing *S. flexneri* infection of the colon; however, since *E. coli* is often only a minor constituent of the human gut microbiota (13), and small animal models fail to recapitulate many aspects of shigellosis in humans (14), the relevance of these data for *S. flexneri* infection in humans is unclear.

Another bacterium found in the gut, *Lactobacillus*, has been of interest to researchers as a probiotic due to its inhibitory effects on enteric pathogens (15). Consistent with the possibility that *Lactobacillus* is protective against *Shigella*, a metagenomic study looking at the association between gut microbiota composition, *Shigella* levels, and diarrheal status in children in low-income countries found that children who were colonized by certain species of *Lactobacillus* had moderate-to-severe diarrhea less often than expected when they were also colonized by *Shigella* (16). This possible role for *Lactobacillus spp*. in preventing symptomatic *S. flexneri* infection is further supported by experiments using cell-culture models that demonstrate that *Lactobacillus spp*. can inhibit *Shigella* attachment to and invasion of colonic epithelial cells (17–19). However, similarly to *E. coli, Lactobacillus spp*. are minor constituents of the gut microbiota (20). Furthermore, only a subset of species of *Lactobacillus* have been observed to stably colonize the intestinal tract, while others are known to only transiently colonize the gut (21). Our understanding of the interactions between *S. flexneri* and prominent members of the gut microbiota and what effects these have on *S. flexneri* virulence remains limited.

The gut microbiota is an incredibly diverse community comprised of hundreds of bacterial species. Most of these bacteria belong to just two phyla, Bacteroidota and Bacillota (formerly called Bacteroidetes and Firmicutes, respectively) (22). While a number of genera from Bacillota can be found in the human gut microbiota, Bacteroidota is largely represented by just four genera, the most abundant of which are *Bacteroides* and *Prevotella* (23). In humans, there appears to be a trade-off between having a *Prevotella* dominated microbiota or a *Bacteroides* dominated one (24–26), with studies suggesting that diet may play a fundamental role in enriching for either *Prevotella*, which is more abundant in people of non-Western societies, or *Bacteroides*, which is more abundant in people of Western societies (27–30). In people from the United States and Western Europe, *Bacteroides* is often the most abundant genus in the gut microbiota (24, 31). Because they are major constituents of the human gut microbiota and are prevalent amongst humans from different populations, *Bacteroides* species have been proposed as model organisms for studying gut microbes (23).

One of the best studied *Bacteroides* species, *Bacteroides thetaiotaomicron* (*Bt*), has been shown to impact the virulence of a number of enteric pathogens (32). *Bt* releases succinate that increases virulence gene expression (33), proteases that cleave the T3SS (34), and fucose that represses T3SS gene expression of enterohemorrhagic *E. coli* (35). Additionally, *Bt* polysaccharides have been shown to inhibit toxin release by *Clostridium difficile* (36). In this study, we examine the effects of *Bt* on *S. flexneri* virulence. We show that *Bt* suppresses *S. flexneri* invasion through the downregulation of *S. flexneri* T3SS gene expression. This suppression is mediated by *Bt* outer membrane vesicles (OMVs), which fuse to *S. flexneri* and post-transcriptionally repress its master virulence regulator, VirF. This demonstrates that a prominent member of the human gut microbiota impacts *Shigella* pathogenesis, and that OMVs can modulate the gene expression of enteric pathogens.

## Results

### *Bacteroides thetaiotaomicron* conditioned medium inhibits *Shigella flexneri* invasion

To determine whether interspecies interactions with members of the human gut microbiota impact *S. flexneri* virulence, we studied the effect that the common gut microbe *B. thetaiotaomicron* (*Bt*) has on *S. flexneri* invasion and cell-to-cell spread (plaque formation) in a monolayer of cultured epithelial cells. We simulated *S. flexneri* encountering an established gut microbiota by growing *S. flexneri* in cell-free conditioned medium (CM) collected from late stationary phase *Bt* cultures. *S. flexneri* grown in brain heart infusion (BHIS) supplemented with *Bt* CM had an 11-fold reduction in invasion rate relative to *S. flexneri* grown in BHIS alone (Fig. 1A). Additionally, *S. flexneri* grown in the presence of *Bt* CM formed fewer plaques than *S. flexneri* grown in BHIS alone (Figs. 1B and 1C), consistent with the lower rate of invasion. Together, this indicates that *Bt* releases a factor that inhibits *S. flexneri* invasion.

**Figure 1:**
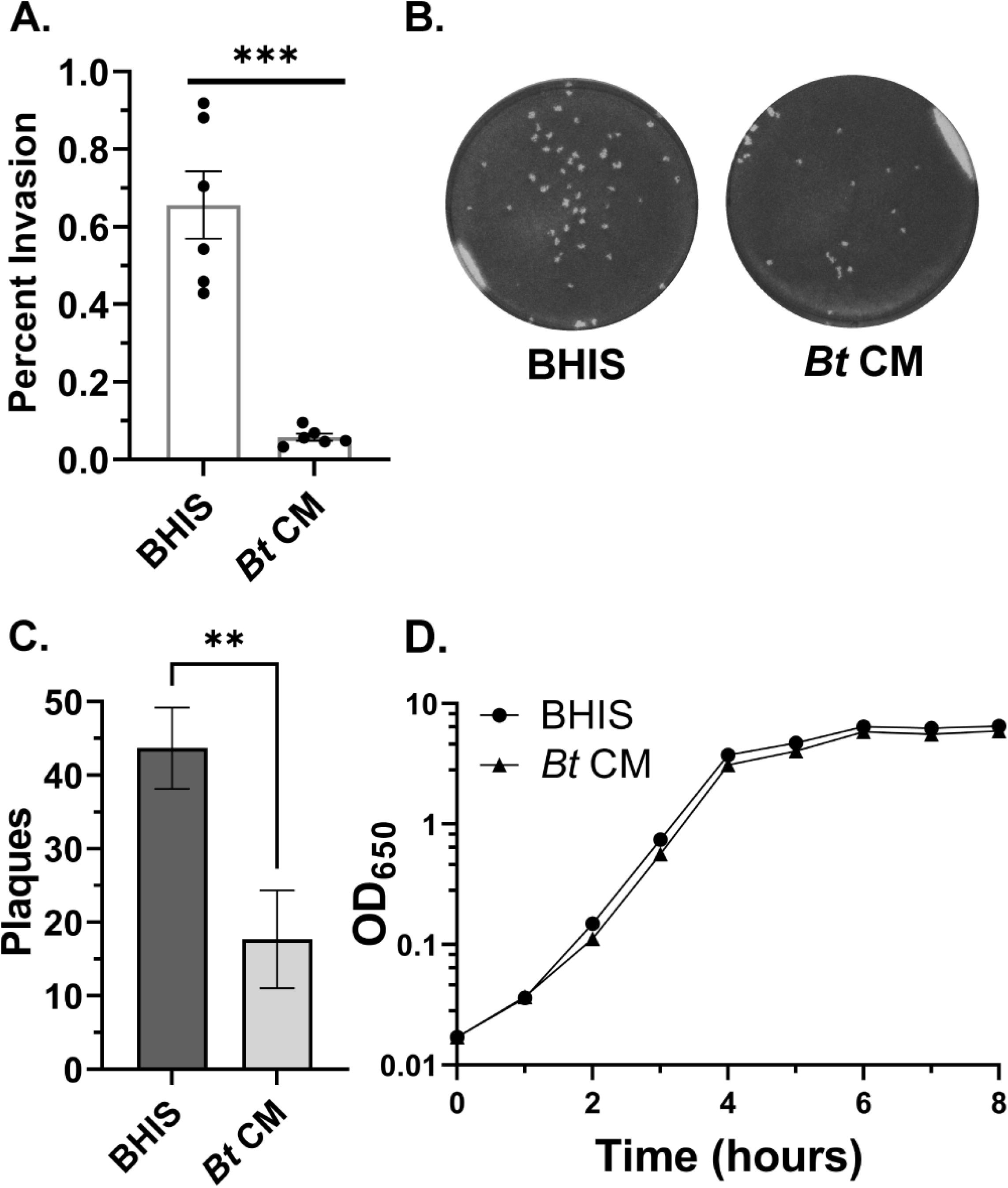
*Bt* CM impacts *S. flexneri* virulence, but not growth, during log phase. Samples labeled *Bt* CM were grown in a mixture of ½ BHIS and ½ *Bt* CM. (A) Invasion rates of *S. flexneri* grown in BHIS or *Bt* CM as measured by a gentamicin protection assay are significantly different (0.66% vs 0.06%, *p* = 0.0008, n = 6). Error bars indicate the standard error of the mean (SEM) calculated from six biological replicates. *P* values were determined by a paired, two-tailed *t* test. (B) A confluent layer of Henle cells was infected with *S. flexneri* grown in BHIS or *Bt* CM. For visualization of plaques, Henle cell monolayers were stained after 72 hours of infection. (C) Average number of plaques formed by *S. flexneri* after growth in BHIS or *Bt* CM was significantly different (43.7 vs 17.7, *p* = .003) as measured from three biological replicates. Error bars indicate standard deviation (SD). *P* values were determined using a paired, two-tailed *t* test. (D) Growth curves of *S. flexneri* in BHIS and *Bt* CM are derived from three biological replicates. *P* values of doubling times were determined using a paired, two-tailed *t* test.

Because it is possible that the observed decrease in invasiveness was due to effects of *Bt* CM on *S. flexneri* growth rate, we measured the growth of *S. flexneri* in the presence and absence of *Bt* CM. The only effect noted on *S. flexneri* growth rate by *Bt* CM was during the first two hours of growth, when the doubling time of *S. flexneri* grown in BHIS supplemented with *Bt* CM (37.6 minutes) was significantly longer than its doubling time in BHIS alone (29.2 minutes, *p* = 0.022, Fig. 1D). However, during mid and late log phase, the growth phase used for the invasion assay, the doubling time of *S. flexneri* in the presence or absence of *Bt* CM was nearly identical (25.7 minutes vs 25.9 minutes, *p* = 0.64, Fig. 1D), indicating that the decrease in *S. flexneri* invasion following growth in *Bt* CM was not due to an effect on growth.

### *Bt* CM reduces the levels of *S. flexneri* virulence factors

Invasion of eukaryotic cells by *S. flexneri* is type three secretion system (T3SS) dependent. Since growth of *S. flexneri* in the presence of *Bt* CM resulted in reduced invasiveness of *S. flexneri* (Fig. 1A), we hypothesized that *Bt* CM represses *S. flexneri* T3SS protein expression. To test this, *S. flexneri* was grown in BHIS supplemented with increasing proportions of *Bt* CM, and the amount of the *S. flexneri* T3SS protein IpaC was determined by Western blot analysis. *Bt* CM inhibited the expression of IpaC in a dose-dependent manner (Fig. 2A). This repression was not unique to IpaC. Other T3SS proteins (IpaA, IpaB, and IpaD) and a non-T3SS associated virulence factor (IcsA) were also produced in lower amounts when *S. flexneri* was grown in the presence of *Bt* conditioned medium (Fig. 2B). In addition to cell-associated virulence factors, secreted virulence factors, including IcsA, IpaA, IpaB, and IpaC, were also decreased in the presence of *Bt* CM relative to BHIS (Fig. S1), suggesting that *Bt* CM reduces the production of *S. flexneri* virulence factors rather than triggers their secretion.

**Figure 2:**
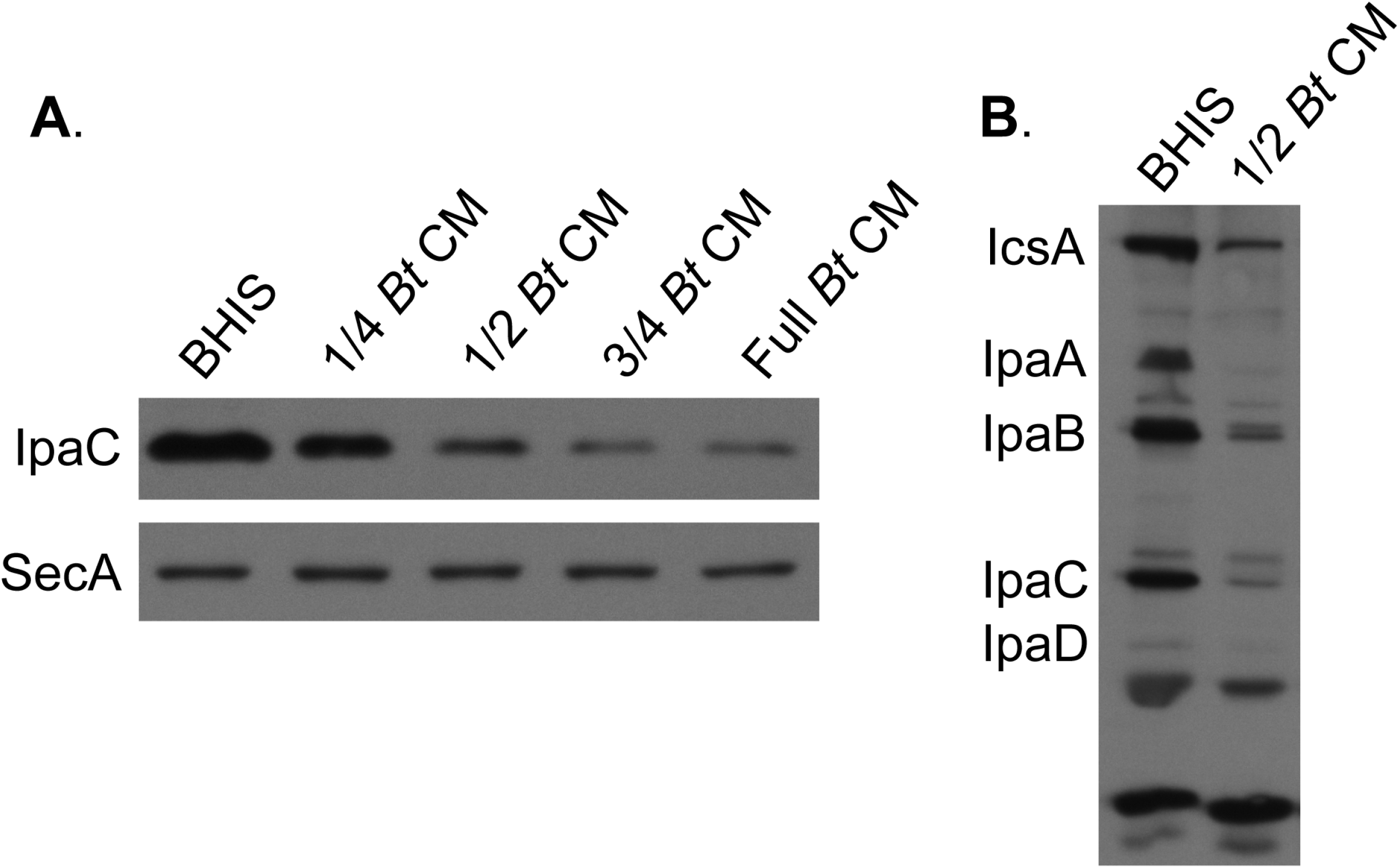
*Bt* CM reduces *S. flexneri* virulence protein levels. (A) Western blot using anti-IpaC antisera was performed on total proteins collected from *S. flexneri* grown to log phase in BHIS supplemented with increasing proportions of *Bt* CM. Anti-SecA antisera was used as a loading control. (B) Western blot using monkey anti-*S. flexneri* convalescent-phase antiserum was performed to look at the protein levels of a panel of *S. flexneri* virulence factors when grown in either BHIS or a mixture of ½ BHIS and ½ *Bt* CM. For both panels, representative Western blots of 3 independent experiments are shown.

### *Bt* CM reduces *S. flexneri* virulence genes at the transcriptional and post-transcriptional level

The inhibitory factor produced by *Bt* could be affecting *S. flexneri* virulence gene expression directly or indirectly by affecting expression of upstream regulators. In *S. flexneri*, VirF is the master virulence transcriptional regulator, which activates expression of a second regulatory gene, *virB*. VirB regulates other virulence genes, including the *ipa* genes. Additionally, *icsA* is directly regulated by VirF, but not by VirB ((6–8), Fig. 3A). Since both IcsA and Ipa protein levels were reduced, this suggested that *Bt* CM represses *virF*. To test whether *Bt* CM affects *S. flexneri* virulence gene expression, the relative virulence gene expression level of *S. flexneri* grown in the presence or absence of *Bt* CM was determined by quantitative reverse transcription PCR (RT-qPCR). *virB* and *icsA* expression were repressed 5.7 and 6.4-fold, respectively, in the presence of *Bt* CM. Additionally, VirB’s downstream targets *ipaB* and *ipaC* were repressed 8.8 and 9.0-fold (Fig. 3B), consistent with *Bt* CM repression of *virB*. Surprisingly, *virF* levels were unchanged. This indicated that either *Bt* CM repression of its two targets, *virB* and *icsA*, was independent of VirF, or that VirF levels were reduced post-transcriptionally. To verify that the effect of *Bt* CM on *S. flexneri* virulence gene expression was specific to *Bt* and not due to general effects of nutrient depletion on *S. flexneri*, CM from *E. coli* MG1655 was collected and its effect on *S. flexneri* virulence gene expression was determined. *E. coli* CM did not affect the expression of any *S. flexneri* virulence genes tested (Fig. S2), indicating that these effects are not simply due to nutrient depletion. To determine if *Bt* CM regulates VirF post-transcriptionally, an S-tag was fused in frame to the 3’ end of *virF*, and the tagged gene was expressed from its native promoter on a plasmid. Western blot analysis of S-tagged VirF showed that VirF protein levels were reduced in *Bt* CM relative to BHIS (Fig. 3C). These data, showing that *Bt* CM reduces VirF S-tag levels (Fig. 3C), but not *virF* gene expression (Fig. 3B), suggest that *Bt* CM affects VirF levels post-transcriptionally.

**Figure 3:**
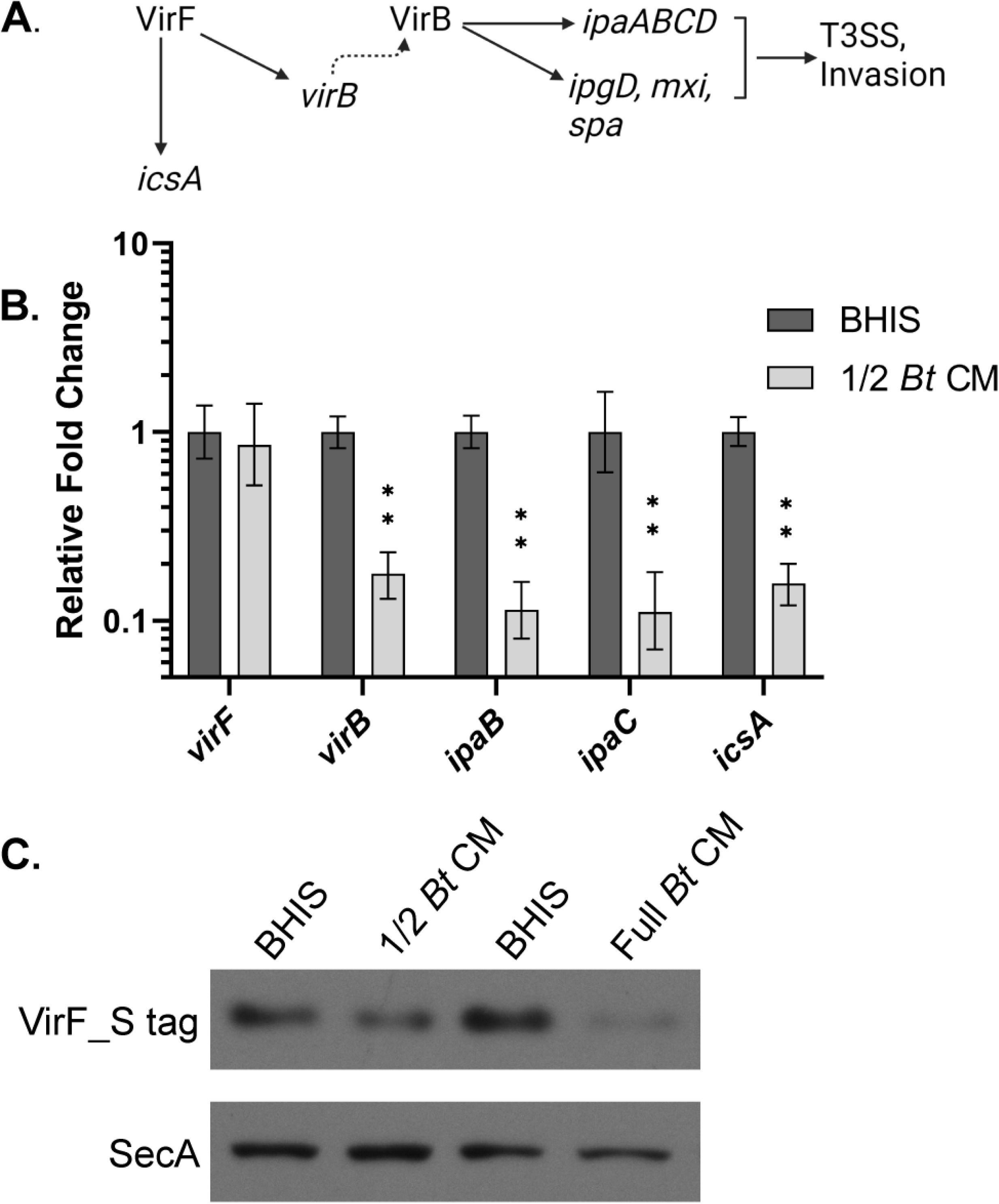
*Bt* CM transcriptionally represses *S. flexneri* virulence gene expression and post-transcriptionally represses the master virulence regulator VirF. (A) Schematic of the virulence gene regulon of *S. flexneri*. (B) Relative virulence gene expression of *S. flexneri* grown to log phase in either BHIS or ½ *Bt* CM was measured by RT-qPCR. *C*_*t*_ values were normalized to the mean of two endogenous controls, *gyrA* and *secA. P* values were determined from the Δ*C*_*t*_ values of three biological replicates using a two-tailed Student’s *t* test with Holm-Šídák correction for multiple comparisons. (**, *p* < 0.01). Error bars indicate SD. (C) Total proteins were collected from *S. flexneri* expressing *virF_S-tag* from the native *virF* promoter on the plasmid pWKS30 grown under the indicated conditions. Western blot was performed on these proteins using an anti-S peptide epitope tag antibody to look at relative VirF_S-tag protein levels. Anti-SecA antisera was used as a loading control. A representative Western blot of more than 3 independent experiments is shown.

### *Bt* inhibitory factor is not a secreted metabolite or protein

To characterize the nature of the *Bt* secreted inhibitory factor, we looked for evidence that the inhibitory factor was either a secreted metabolite or protein. Bacteria of the gut microbiota, including *Bacteroides* species, are known to excrete a variety of short chain fatty acids (SCFAs) (37, 38), and enteric pathogens have been shown to modulate their virulence genes in response to exogenous SCFAs (39–41). To determine whether *S. flexneri* was responding to a secreted metabolite, such as a SCFA, the small molecules (< 10 kDa) were separated from proteins and other large molecules using a protein concentrator (Fig. 4A). The inhibitory activity of fractionated *Bt* CM was associated with the retentate; the small molecules in the flow-through had no effect on the virulence protein levels (Fig. 4B, lanes 5 and 6). We repeated the fractionation of *Bt* CM with a 100 kDa molecular weight cutoff filter. Similar to what was observed with the 10 kDa cutoff filter, the majority of the inhibitory activity remained in the retentate (Fig. 4B, lanes 9 and 10), suggesting that the active factor was not a small molecule. As expected, fractionating BHIS with the protein concentrators had no effect on IpaC expression relative to unfractionated BHIS (Fig. 4B, lanes 3-4 and 7-8). The lack of effect of the flow-through on the level of *S. flexneri* virulence proteins indicates that the inhibition is due to the presence of a high-molecular weight factor and further demonstrates that it is not a result of depletion of a growth factor from the conditioned medium.

**Figure 4:**
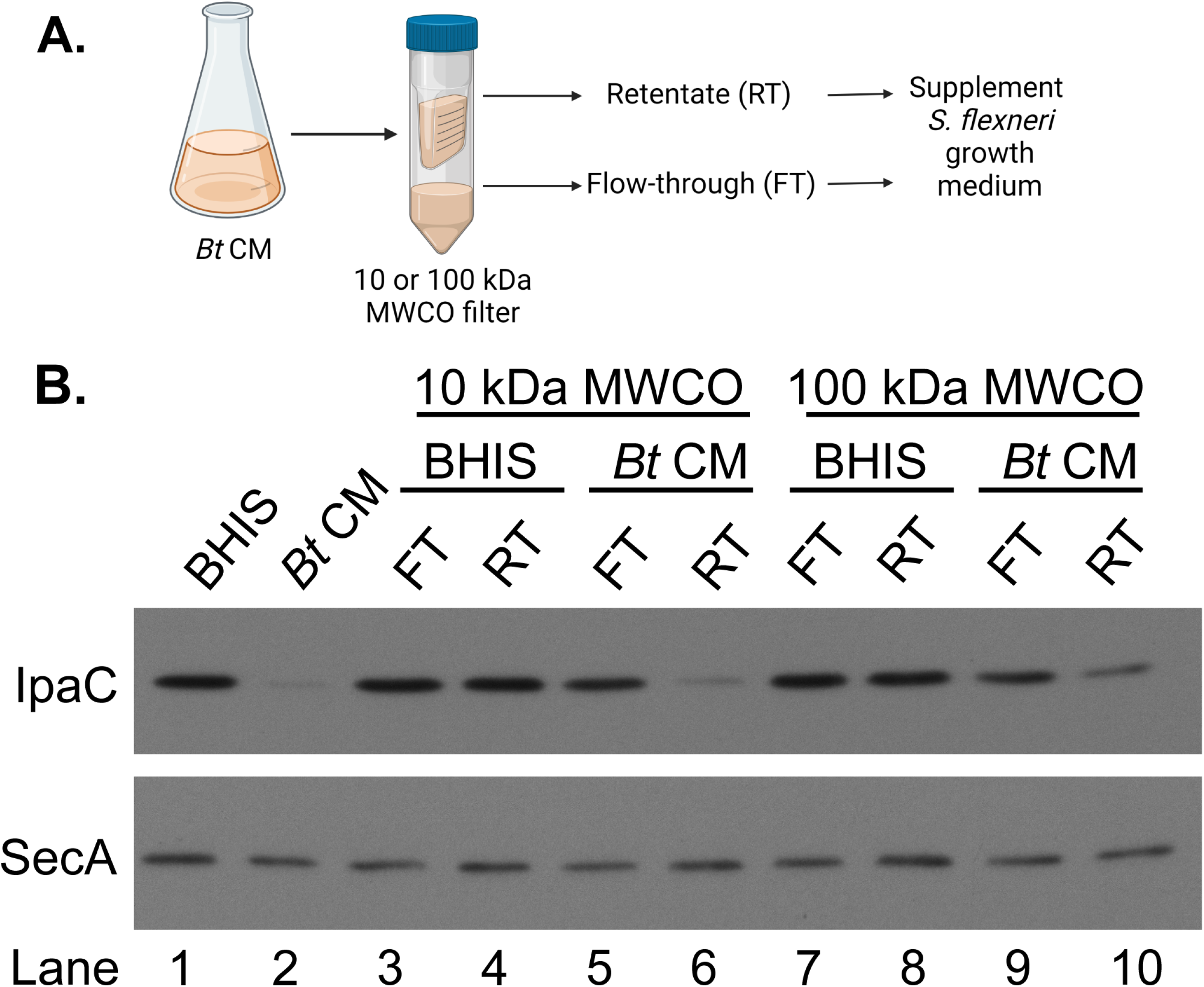
Fractionation of *Bt* CM shows that the active fraction is not a small molecule. (A) *Bt* CM was fractionated by size using either a 10 kDa molecular weight cutoff (MWCO) filter or a 100 kDa MWCO filter. The retentate fraction (RT) or the flow-through (FT) was added to *S. flexneri* growth medium. As a negative control, BHIS was subjected to the same fractionation. (B) Growth in the presence of the retentate fraction of *Bt* CM from either a 10 kDa or 100 kDa filter inhibits *S. flexneri* IpaC expression. Anti-SecA was used as a loading control. A representative Western blot of 3 independent fractionation experiments is shown.

To assess whether the inhibitor was a protein, *Bt* CM was treated with Proteinase K. However, Proteinase K treatment of *Bt* CM did not eliminate its ability to inhibit *S. flexneri* IpaC production (Fig. 5A), suggesting the active component was not a protein. It was possible that the *Bt* secreted factor was a soluble metabolite that non-specifically stuck to the protein concentrator during fractionation and was recovered in the retentate fraction. To rule this out, we fractionated *Bt* CM by ultracentrifugation. This separated the supernatant, containing soluble molecules, from a pellet, containing the lipids and other insoluble components of the CM. These two fractions were then tested individually for their effect on *S. flexneri*, where it was observed that *S. flexneri* grown in medium containing the resuspended pellet had reduced IpaC expression (Fig. 5B, lane 6), while *S. flexneri* grown in medium supplemented with the supernatant (lane 5) had IpaC levels comparable to the BHIS control (lane 1) (Fig. 5B). This indicated that the active component in *Bt* conditioned medium was insoluble in aqueous solution and likely to be lipid associated. To ensure that the inhibitory factor being ultracentrifuged out of *Bt* CM was specific to the CM, BHIS was fractionated in the same way. Neither the supernatant nor pellet isolated from BHIS had any effect on *S. flexneri* IpaC levels (Fig. 5B, lanes 1-3).

**Figure 5:**
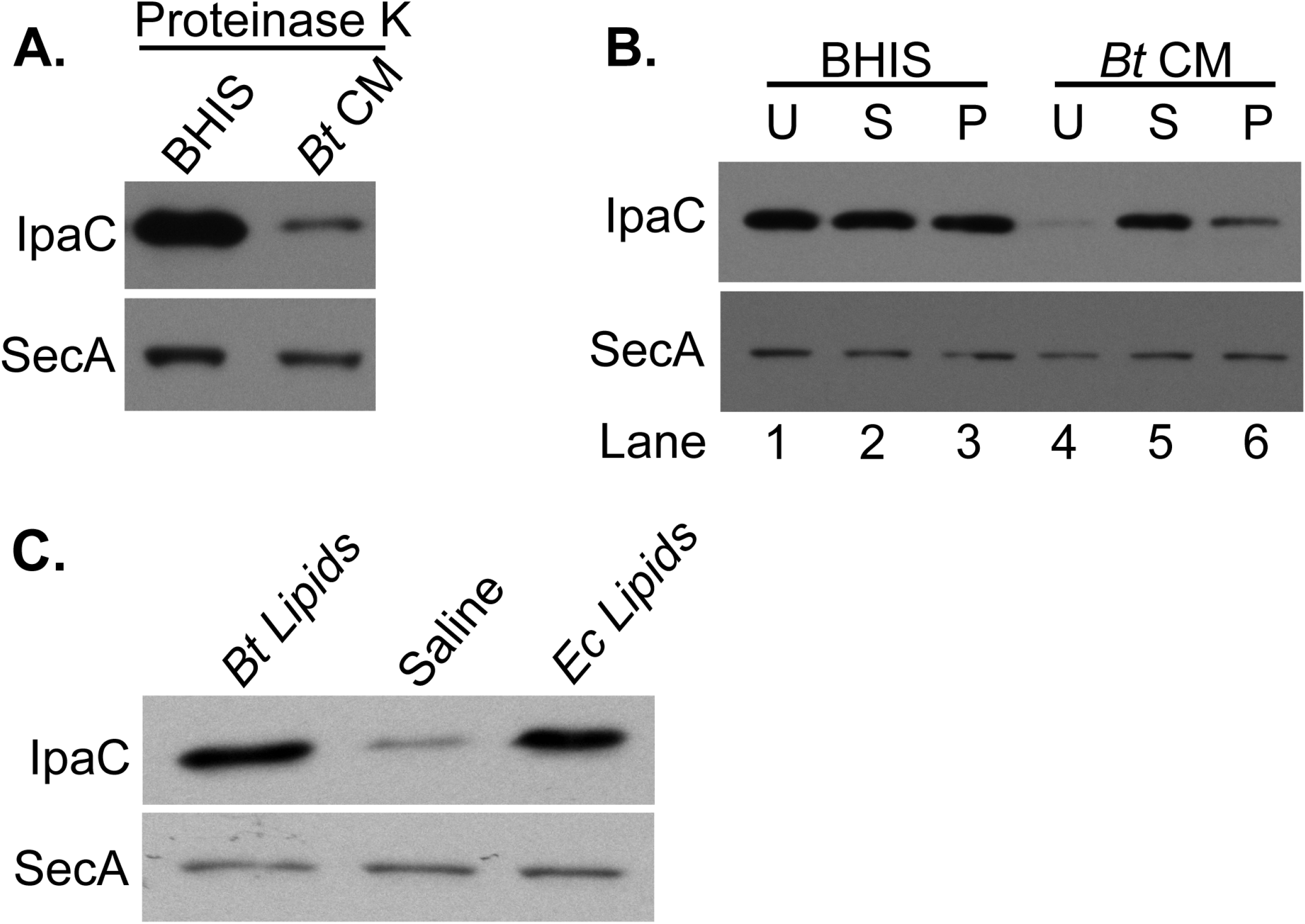
Active component of *Bt* CM is lipid-associated and Proteinase K resistant. (A) To degrade proteins, BHIS and *Bt* CM were treated with 50 µg/mL of Proteinase K. Each was then mixed 1:1 with untreated BHIS. Total proteins were collected from *S. flexneri* grown to mid log phase in either Proteinase K treated BHIS or Proteinase K treated *Bt* CM, and used for an anti-IpaC Western blot. (B) BHIS and *Bt* CM were ultracentrifuged to separate the water-soluble supernatant from the insoluble pellet. The supernatant was mixed 1:1 with BHIS, while the pellet was resuspended in BHIS. Total proteins were collected from *S. flexneri* grown to log phase in unfractionated medium (U), or BHIS supplemented with either the supernatant (S) or pellet (P) fractions and used for an anti-IpaC Western blot. (C) *S. flexneri* was grown in the presence of *Bt* lipids, *E. coli* (*Ec*) lipids, or an equal volume of saline. The effect of these lipids on *S. flexneri* IpaC levels was determined by Western blot. For all three panels, anti-SecA was used as a loading control. Each fractionation or isolation and their corresponding Western blot were performed at least 3 independent times.

### *Bt* lipids and OMVs inhibit *S. flexneri* virulence factors

To directly determine whether the inhibition of *S. flexneri* virulence factor production was due to the presence of *Bt* lipids in the culture supernatant, we extracted the total lipids from a pellet of stationary phase *Bt* and grew *S. flexneri* in the presence of these lipids. Growth in the presence of total *Bt* lipids repressed IpaC expression (Fig. 5C). This effect was specific to *Bt* lipids, since an equivalent amount of lipids from *E. coli* had no effect (Fig. 5C).

Gram-negative bacteria, including *Bt*, are known to produce outer membrane vesicles (OMVs). OMVs have been shown to be involved in cell-to-cell communication (42). Because the size and lipid nature of the inhibitory factor was consistent with OMVs, we purified OMVs from the *Bt* CM for analysis. Transmission electron microscopy confirmed the presence of small spherical structures (Fig. 6A) ranging from 20 nm to 100 nm in diameter, with an average diameter of 51 nm (Fig. 6B). These structures were similar in both size and appearance to OMVs previously isolated from other *Bacteroides* species (43). Additionally, we validated that the vesicles isolated from *Bt* CM were OMVs by utilizing *Bt* strains expressing tagged proteins known to localize to either the outer membrane (OM) or OMVs extracted from these strains. Consistent with the results of Valguarnera *et al*. (44), BT_1488 was localized to the isolated extracellular vesicles, while BT_0418 was found in the OM and not in the extracellular vesicles (Fig. S3). This indicates that the purified fraction contained *Bt* OMVs and was not significantly contaminated with cellular membranes.

**Figure 6:**
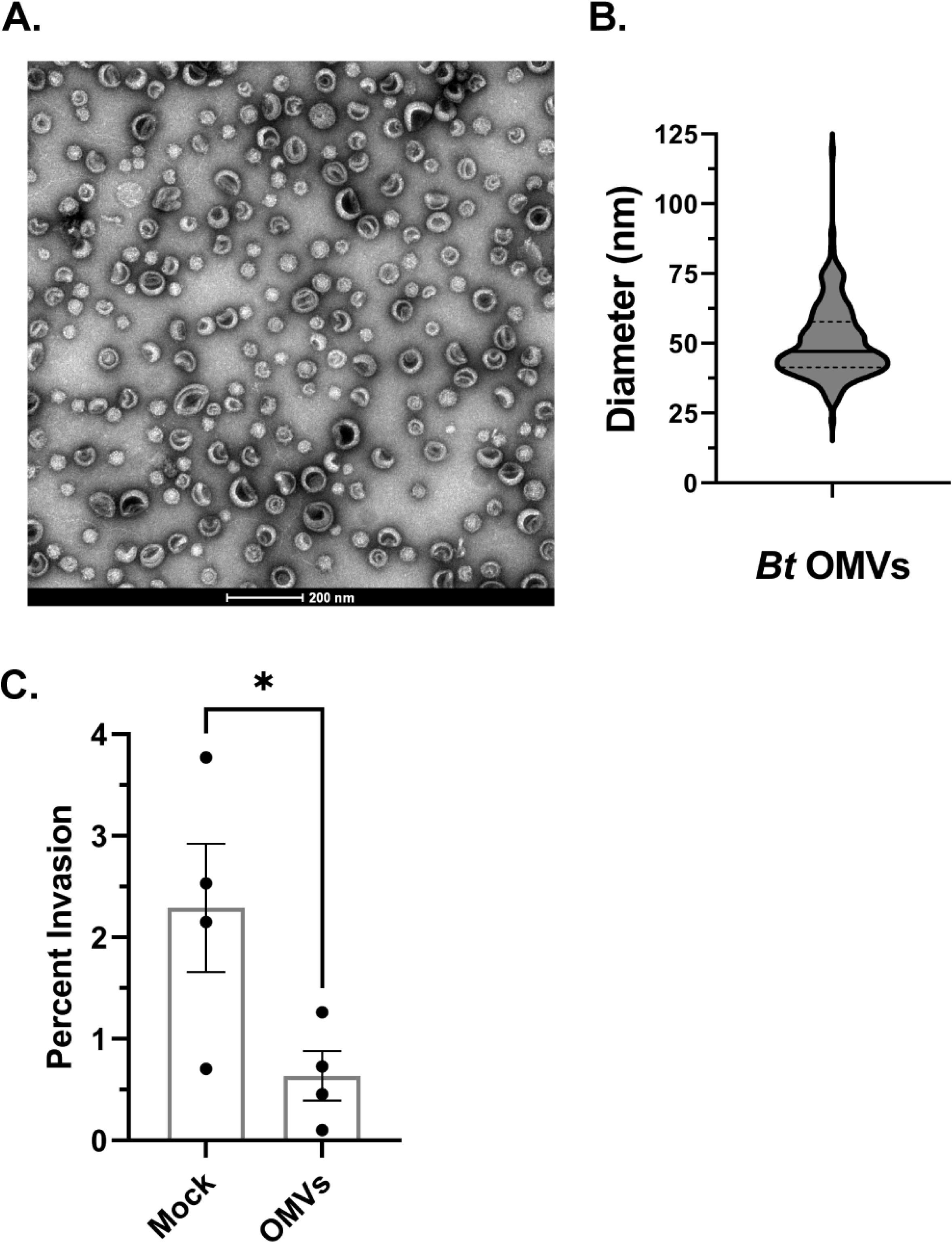
*Bt* CM contains OMVs that inhibit *S. flexneri* invasion. (A) Extracellular vesicles were concentrated from *Bt* CM, purified on a density gradient, and counter stained with 2% uranyl acetate for visualization by transmission electron microscopy. A representative image taken at 60,000x magnification is shown. Scale bar = 200 nm. (B) The diameters of 400 OMVs were measured using ImageJ (91). (C) A gentamicin protection assay was used to determine the invasion rate of *S. flexneri* grown in the presence of 20 µg/mL of *Bt* OMVs or an equal volume of mock extract. *P* value was determined by a two-tailed, unpaired *t* test from 4 biological replicates (*, *p* < 0.05). Error bars indicate SEM.

To determine whether *Bt* OMVs are the factor responsible for repressing *S. flexneri* virulence in *Bt* CM, the invasion rate of *S. flexneri* grown in the presence of *Bt* OMVs was measured. The amount of OMVs added was normalized to protein levels in the purified vesicles and is consistent with concentrations used in previous studies (45, 46). We observed that when treated with *Bt* OMVs, *S. flexneri* was less invasive than *S. flexneri* treated with an extract of uninoculated BHIS (mock extract) (Fig. 6C). Similar to the results with *Bt* CM, *Bt* OMVs inhibited expression of *S. flexneri* IpaC (Fig. 7A) and the virulence factors IcsA, IpaA, and IpaB (Fig. 7B). As determined by RT-qPCR, this reduction in protein levels was associated with reduced expression of the virulence genes *virB, icsA, ipaB*, and *ipaC*. Relative to growth in BHIS alone, *S. flexneri* grown in the presence of *Bt* OMVs had reductions in expression of 3.9-fold for *virB*, 3.5-fold for *icsA*, 5.8-fold for *ipaB*, and 5.7-fold for *ipaC*. Also consistent with the results from *Bt* CM treatment, v*irF* gene expression was unchanged by growth in the presence of the *Bt* OMVs (Fig. 7C), but protein levels of VirF were reduced (Fig. 7D). Thus, the gene and protein expression patterns observed in *Bt* CM were recapitulated by *Bt* OMVs. The level of repression of the virulence genes by the vesicles is somewhat less than that observed with the conditioned medium. This may reflect differences in the amount of the inhibitory factor between the purified vesicles and the crude conditioned medium, or there may be an additive effect of the vesicles and some other component in the conditioned medium.

**Figure 7:**
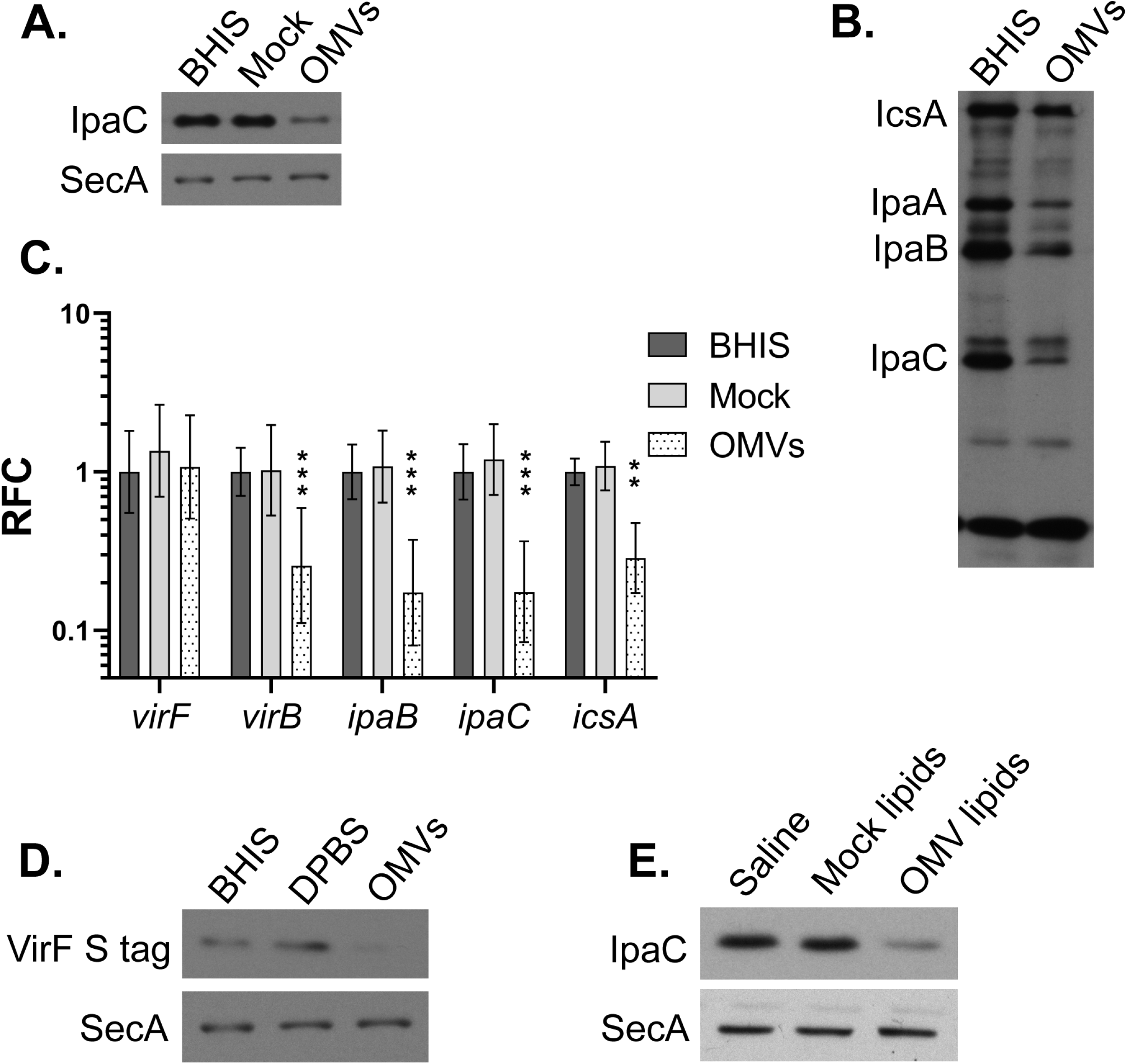
*Bt* OMVs repress *S. flexneri* virulence gene expression. (A) An anti-IpaC western blot was performed on proteins collected from *S. flexneri* that was either untreated, treated with mock extract, or treated with 20 µg/mL *Bt* OMVs and grown to log phase. Anti-SecA was used as a loading control. (B) Proteins were collected from untreated *S. flexneri* and *S. flexneri* treated with 20 µg/mL of *Bt* OMVs. A Western blot using monkey anti-*Shigella* convalescent-phase antiserum was performed to look at the relative protein levels of a panel of virulence factors. (C) Relative fold change (RFC) of virulence gene expression of *S. flexneri* treated with either mock extract or 20 µg/mL of *Bt* OMVs compared to untreated *S. flexneri* was measured by RT-qPCR. *C*_*t*_ values were normalized to the mean of two endogenous controls, *gyrA* and *secA. P* values were determined from the Δ*C*_*t*_ values of three biological replicates using two-way ANOVA with Dunnett’s posttest for multiple comparisons of gene expression in treated samples to the untreated samples. (*, *p* < 0.05, **, *p* < 0.01, ***, *p* < 0.001). Error bars indicate standard deviation (SD). (D) An anti-S peptide Western blot was performed on proteins collected from *S. flexneri* pWKS30::*virF*_*S-tag* that was either untreated, treated with Dulbecco’s phosphate buffered saline supplemented with salts (DPBS), or treated with 40 µg/mL *Bt* OMVs and grown to log phase. Anti-SecA was used as a loading control. (E) *S. flexneri* was grown to log phase in the presence of saline, lipids extracted from *Bt* OMVs, or an equal volume of mock lipids. An anti-IpaC Western blot was performed. Anti-SecA was used as the loading control. Each Western blot in this figure is representative of at least 3 independent replicates, except for panel B, which is representative of 2 independent replicates.

### *Bt* OMV lipids inhibit IpaC

Since *Bt* lipids and OMVs both repress *S. flexneri* IpaC levels, we wanted to determine whether *Bt* OMV lipids alone were sufficient to inhibit *S. flexneri* IpaC expression. Lipids extracted from purified *Bt* OMVs repressed *S. flexneri* IpaC levels (Fig. 7E), indicating that the inhibitory effect of the OMVs is due to the OMV lipids themselves and not to the presence of an unknown cargo contained in the *Bt* OMVs. Performing the same OMV and lipid extraction starting with BHIS (mock lipids), and growing *S. flexneri* in the presence of this extract had no effect on IpaC levels (Fig. 7E), demonstrating that the extracted lipids that inhibit *S. flexneri* virulence protein levels are specific to *Bt* OMVs and not present in the uninoculated BHIS media.

### *Bt* OMVs directly interact with *S. flexneri*

Because the effect of *Bt* produced OMVs on *S. flexneri* invasion is due to the lipids in *Bt* OMVs, we hypothesized that the OMVs were fusing with *S. flexneri*’s membrane, and the presence of *Bt* lipids in the membrane of *S. flexneri* could initiate the observed changes in *S. flexneri* virulence protein levels. To test whether the *Bt* vesicles directly contact *S. flexneri, Bt* OMVs were stained with the lipophilic fluorescent dye FM4-64 FX and washed extensively to remove unbound dye. The stained *Bt* OMVs were then co-incubated with *S. flexneri*, and the amount of fluorescence associated with the *S. flexneri* cells was determined over time. After 30 minutes of coincubation, *S. flexneri* exhibited a fluorescent signal of 24.2 RFU/OD_650_, and after 2 hours of coincubation, the fluorescence of *S. flexneri* increased to 36.6 RFU/OD_650_ (Fig. 8). This suggested that the fluorescently labeled OMVs fused with *S. flexneri*. Since free FM4-64 FX can directly bind to *S. flexneri*, we controlled for free dye carryover by performing the same staining and washing procedure on saline that did not contain OMVs. After 30 minutes and 2 hours of coincubation with the stained and washed control saline, *S. flexneri* only exhibited fluorescence of 1.1 and 1.3 RFU/OD_650,_ respectively. This ruled out dye carryover from the staining and washing procedures as the cause of *S. flexneri* fluorescence. To further control for release of dye from the vesicles during co-incubation, labelled vesicles were soaked in buffer for the same amount of time used in the transfer experiment. After soaking, the vesicles were removed by ultracentrifugation, and *S. flexneri* was incubated with either the soaked OMVs or the buffer in which the OMVs had been soaked. While some fluorescence was measured in *S. flexneri* that was resuspended in the soaking buffer, the amount of fluorescence was over 10 times lower than that of *S. flexneri* that was incubated with the soaked OMVs. Additionally, the fluorescence of *S. flexneri* that was co-incubated with soaked OMVs was indistinguishable from that of *S. flexneri* that was co-incubated with unsoaked OMVs (Fig. S4). Together, this suggests that passive transfer of dye from the OMVs to *S. flexneri* via diffusion of dye away from the stained vesicles is only a minor contributor to *S. flexneri* fluorescence and indicates that fusion or direct contact between the OMVs and *S. flexneri* is likely occurring.

**Figure 8:**
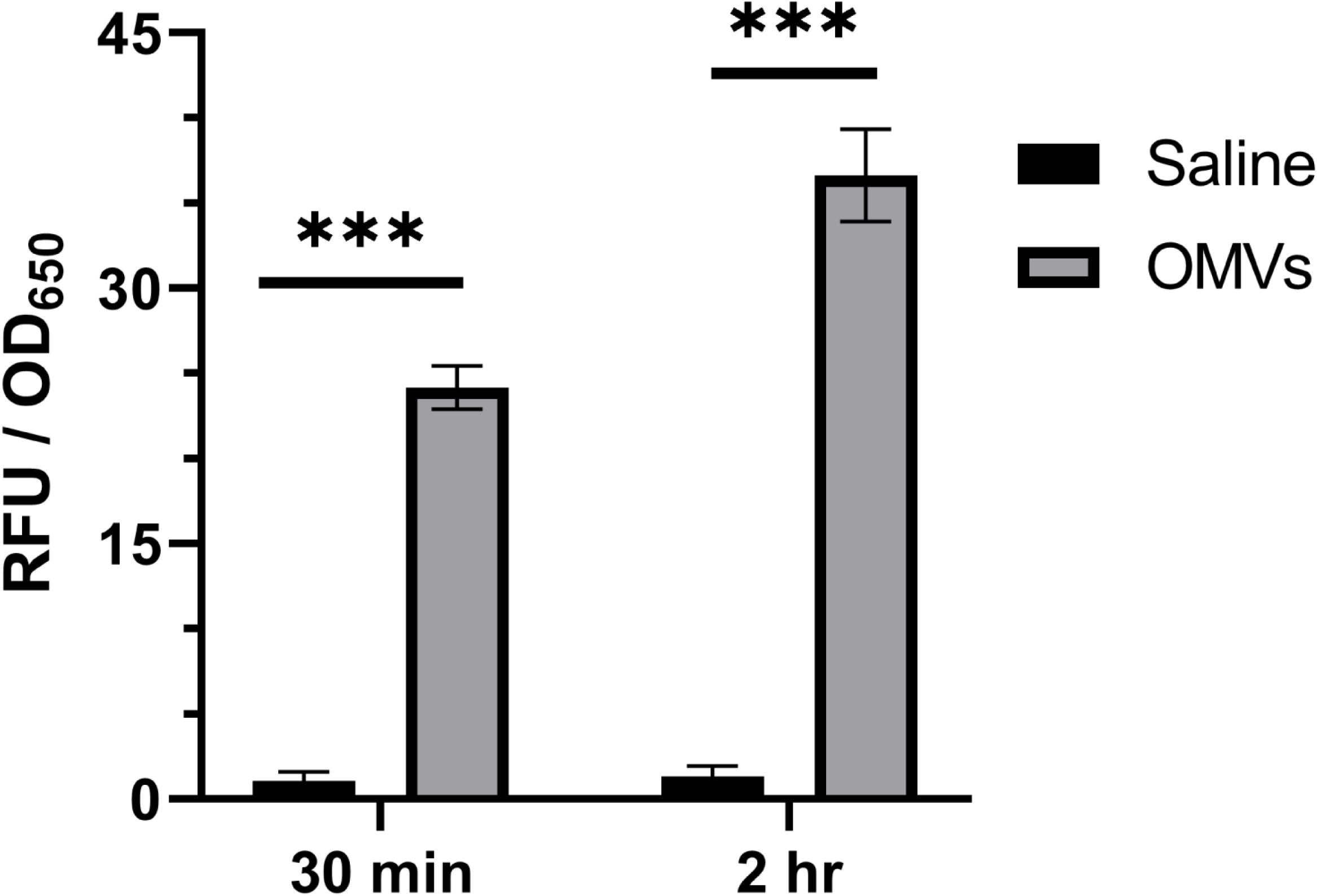
*Bt* OMVs fuse to *S. flexneri*. Saline or *Bt* OMVs were stained with 5 µg/mL of the fluorescent dye FM4-64 FX, washed extensively, and incubated with *S. flexneri* for 2 hours at room temperature. At timepoints of 30 minutes and 2 hours, *S. flexneri* was fixed with 4% PFA and its fluorescence measured on a plate reader (Ex/Em of 510/640 nm). *P* values were determined from three biological replicates using a two-tailed Student’s *t* test with Holm-Šídák correction for multiple comparisons. (***, *p* < 0.001). Error bars indicate standard deviation.

## Discussion

Despite the fact that *S. flexneri* must interact with the gut microbiota before establishing infection in the colon, the impact of these interactions on *S. flexneri* pathogenesis remains poorly understood. In this study, we show that the common gut microbe *B. thetaiotaomicron* produces OMVs that repress *S. flexneri* virulence gene expression and inhibit *S. flexneri* invasion.

These results have implications in the broader context of *S. flexneri* pathogenesis. First, it is possible that gut microbiotas enriched for *Bt* have a protective effect, making the people that harbor these microbiotas more resistant to *S. flexneri* infection. In humans, *Bacteroides* is often the most abundant genus in the gut microbiota. However, it also the most variable, ranging from comprising the majority of gut microbes in some people to a small minority in others (22, 24). Our data show that the inhibitory effect of *Bt* on *S. flexneri* virulence protein levels is dose dependent, with a higher proportion of *Bt* CM leading to a stronger reduction in the T3SS protein IpaC (Fig. 2A), and that the presence of OMVs in *Bt* CM is responsible for its inhibitory effect on *S. flexneri* invasion (Fig. 6). We speculate that if the abundance of *Bt* in the gut microbiota were to correlate with the accumulation of more *Bt* OMVs in the lumen of the colon, then it is conceivable that individuals with higher proportions of *Bt* could be conferred some degree of protection from *Shigella* infection. However, existing data suggesting *Bt* plays a protective role in *S. flexneri* infection are scarce. One study examined the composition of the gut microbiota in children in low-income countries in the presence or absence of *Shigella* (16). These data show that the relative abundance of *Bacteroides* is lower in children who are colonized by *Shigella* and have diarrhea than in children who are colonized by *Shigella*, but don’t have diarrhea. While this correlation between *Bacteroides* proportion and disease state is consistent with the possibility that *Bacteroides* plays a protective role in *Shigella* infection, it is worth noting that a similar decrease in abundance of *Bacteroides* was observed in children who had diarrhea relative to children who did not have diarrhea, even when these children did not have *Shigella*. Thus, it is difficult to parse out whether these effects are due to interactions between *Bacteroides* and *Shigella*, or whether they are simply due to effects of diarrhea on gut microbiota composition. More research is needed to determine whether the abundance of certain species in the gut microbiota can affect susceptibility to *S. flexneri* infection.

Alternatively, it is possible that, rather than repressing *S. flexneri* virulence, *Bt* functions to help coordinate *S. flexneri* invasion. The low infectious dose of *Shigella* (47) suggests that its invasion process is both highly coordinated and efficient. Furthermore, a number of cues, including temperature (48), pH (49), oxygen tension (50), bile salts (51), and osmolarity (52), that help coordinate *S. flexneri* invasion have been identified. Perhaps *Bt* produced OMVs are functioning as another cue that prevents *S. flexneri* from prematurely expressing its T3SS in the lumen of the colon, before other cues closer to the colonic epithelium trigger T3SS expression at the appropriate time. This paradigm, where members of the gut microbiota secrete cues that are sensed by enteric pathogens and used to coordinate infection, has been reported in other enteric pathogens (33, 35).

The inhibitory effect of *Bt* CM on *S. flexneri* virulence is due to the presence of *Bt* OMVs (Figs. 6-7). OMVs derived from commensal bacteria are known to modulate the intestinal immune response, deliver molecules that protect mice from colitis, and function in inter-kingdom signaling between bacteria and intestinal cells, while OMVs from pathogenic bacteria have been shown to deliver virulence factors to intestinal cells (53–58). In the context of inter-bacterial interaction, OMVs have been observed to affect population and community dynamics by functioning in both intraspecies and interspecies signaling (42, 59, 60). In this study, we add to the variety of ways that OMVs are known to function in interspecies interaction by showing that OMVs secreted by one species of bacteria can modulate the virulence gene expression of another species. Additionally, by showing that gut microbiota-derived OMVs can modulate the invasiveness of an enteric pathogen, we have identified a possible new function for bacterial OMVs in the colon. Whether this inhibitory effect is specific to *Bt* OMVs and *S. flexneri* virulence genes or is more generalizable to other combinations of gut microbes and enteric pathogens is unknown.

The mechanism by which *Bt* OMVs repress *S. flexneri* virulence gene expression remains to be determined. The interaction between the bacteria involves direct contact between *Bt* OMVs with *S. flexneri*, the majority of which occurs within the first 30 minutes of coincubation (Fig. 8). This suggests that the vesicles are fusing with the *S. flexneri* cells. Additionally, incubation with *Bt* OMVs leads to a decrease in protein levels of *S. flexneri’s* master virulence regulator, VirF (Fig. 7D). The link between OMV fusion and repression of VirF remains uncharacterized; however, it is possible that the fusion of lipids contained in *Bt* OMVs into the membrane of *S. flexneri* triggers stress responses. Virulence gene expression in *S. flexneri* and other enteric pathogens is known to be closely intertwined with bacterial stress responses (61–66). *Bt* OMVs fusing to *S. flexneri* may activate envelope stress response two-component systems, and then these directly or indirectly transduce the signal of *Bt* OMV fusion from the outer membrane of *S. flexneri* to VirF. While *Bt* OMV-induced repression of VirF protein was observed, no change in *virF* RNA was detected (Fig. 7C). This suggests post-transcriptional regulation or effects on the stability of VirF. To date, two examples of post-transcriptional regulation of VirF have been described. Specifically, deletion of two genes involved in tRNA modification, *tgt* and *miaA*, have been shown to decrease the translation efficiency of *virF* despite having no effect on its transcription (7, 67, 68). Future studies will be aimed at determining how OMV fusion leads to the observed decrease in VirF.

Our data suggests that the particular component of *Bt* OMVs that causes the repression of *S. flexneri* virulence gene expression is a lipid. Total lipid extracts from both *Bt* stationary phase cell pellet and *Bt* OMVs repress *S. flexneri* IpaC protein expression, while an equivalent amount of lipids from *E. coli* does not (Fig. 5C). The particular *Bt* lipid or lipids that are responsible for this effect remains unknown. Since *E. coli* lipids do not induce the same effect, it does not appear to be a general lipid effect that reduces IpaC levels. Furthermore, this specificity for *Bt* lipids indicates that the responsible inhibitory lipid(s) are likely enriched in *Bt* relative to *E. coli*, or absent from *E. coli* altogether. *Bt* membrane sphingolipids are a possible candidate. Unlike *E. coli* and almost all other bacteria, members of the phylum *Bacteroidota* (including *Bt*) contain membrane sphingolipids, which comprise over 50% of the total lipids in *Bt* OMVs (69–71). Sphingolipids of both mammalian and bacterial origin can be found in the colon, where *Bt* sphingolipids are known to promote intestinal homeostasis (72, 73). In the environment, the bacterium *Algoriphagus machipongonensis* induces multicellular rosette formation of the choanoflagellate *Salpingoeca rosetta* by releasing OMVs that fuse to *S. rosetta*. The active signal contained in the OMVs that induce *S. rosetta* rosette formation are sulfonolipids, a structural analogue of sphingolipids (74, 75). A similar signaling mechanism may be occurring in *S. flexneri*, whereby *Bt* sphingolipids, released as part of OMVs, fuse to *S. flexneri* and cause the observed repression of virulence gene expression.

## Methods

Bacterial strains and growth conditions: Bacterial strains and plasmids used in this study can be found in Table S1. *S. flexneri* and *E. coli* strains were maintained at -80°C in tryptic soy broth (TSB) containing 20% (vol/vol) glycerol. *S. flexneri* was grown aerobically at 37°C on TSB agar with 0.01% Congo red dye. Overnight cultures in TSB were sub-cultured 1:100 into the indicated growth medium and grown aerobically (200 RPM, 37°C). *E. coli* strains were grown on Luria-Bertani (LB) agar (1% Tryptone, 0.5% yeast extract, 1% NaCl, and 1% agar) and in LB broth (1% Tryptone, 0.5% yeast extract, 1% NaCl) under the same growth conditions as *S. flexneri*.

*B. thetaiotaomicron* (*Bt*) strains were maintained at -80°C in brain heart infusion (BHI, Bacto) containing 5 mg/L hemin and 20% glycerol [vol/vol]. *Bt* was grown on BHI agar supplemented with yeast extract (0.5% [wt/vol]), sodium bicarbonate (0.2% [wt/vol]), 5 mg/L hemin, and L-cysteine (free base, 0.1% [wt/vol]). For liquid culture, *Bt* was grown in supplemented brain heart infusion (BHIS) containing 37 g/L brain heart infusion (Bacto), yeast extract (0.5% [wt/vol]), 5 mg/L hemin, L-cysteine (free base, 0.1% [wt/vol], and 50 mM HEPES sodium salt (pH 7.5). *Bt* was cultured in an anaerobic chamber (Coy) using an atmosphere of 85% N_2_, 10% CO_2_, 5% H_2_ at 37°C. Antibiotics were used at the following concentrations: gentamicin (20 µg/mL) and erythromycin (25 µg/mL).

Construction of plasmids: C-terminal S-tagged (76) VirF (VirF_S-tag) was constructed using the primers indicated in Table S2. The promoter region and coding sequence of *virF* were amplified with the primers from the virulence plasmid of *S. flexneri* 2457T along with the S tag, which was included in the sequence of the reverse primer. The region directly downstream to *virF* on the virulence plasmid was amplified as well. These two pieces were attached using splice by overlap extension PCR (77) and then ligated into the EcoRI and SalI sites of the low-copy vector pWKS30 (78).

Cell culture media and growth conditions: Minimal essential medium (MEM, Gibco) containing heat-inactivated fetal bovine serum (Gibco, 10%, [vol/vol]), tryptone phosphate broth (Bacto, 10%, [wt/vol]), 1X nonessential amino acids (Gibco), and 2 mM glutamine was used to grow Henle cells (intestine 407; ATCC CCL-6). Henle cells were incubated at 37°C with 5% CO_2_.

*Bt* cell free conditioned medium (CM): To generate *Bt* CM, *Bt* was grown anaerobically on BHI agar plus gentamicin. BHIS was inoculated with a single colony of *Bt* and grown anaerobically for 40 hours to late stationary phase. The stationary phase culture was centrifuged at 13,000 x *g* for 10 minutes. The supernatant was filter sterilized using a 0.22 µm PES filter, pH-adjusted to 6.8 with 1M NaOH, and then passed through a second 0.22 µm filter.

Virulence assays: *S. flexneri* invasion was measured by a gentamicin protection assay (79). *S. flexneri* was grown to late log phase in the indicated growth medium. Approximately 10^8^ *S. flexneri* (multiplicity of infection ∼ 100) were added to a confluent monolayer of Henle cells in a 35 mm, 6-well polystyrene plate (Corning) and centrifuged at 1,000 x *g* for 10 minutes. The plate was incubated for 30 minutes at 37°C/5% CO_2_, after which each well was washed 4 times with phosphate-buffered saline (PBS-D) (1.98 g/L KCl, 8 g/L NaCl, 0.02 g/L KH_2_PO_4_, 1.39 g/L K_2_HPO_4_), filled with MEM containing 20 µg/mL gentamicin, and incubated for another hour. The wells were then washed twice more with PBS-D and lysed at room temperature for 10 minutes with 0.1% Triton X-100 to recover intracellular bacteria. Ten-fold serial dilutions of both the input and output bacteria were plated, and invasion efficiency for each sample was calculated as percent invasion (output CFU/input CFU). For plaque assays, *S. flexneri* was grown to log phase in LB, 5 × 10^4^ bacteria were added to a confluent layer of Henle cells in a 6-well polystyrene plate, and centrifuged at 1,000 x *g* for 10 minutes. The plate was incubated for 30 minutes, after which each well was washed 4 times with PBS-D, and the media was replaced with MEM containing 0.45% glucose and gentamicin for 24 hours. After 24 hours, the media was replaced with MEM containing only gentamicin and the plate was incubated for 48 hours more. The wells were washed with PBS-D, fixed with 80% methanol for 5 minutes, and then stained with 0.5% crystal violet for visualization.

SDS-PAGE and immunoblotting: *S. flexneri* was grown to late log phase in the indicated growth medium, centrifuged, resuspended in Laemmli SDS sample buffer (5% β-mercaptoethanol, 3% [wt/vol] SDS, 10% glycerol, 0.02% [wt/vol] bromophenol blue, 63 mM Tris-Cl [pH 6.8]) (80) at a concentration of approximately 2 × 10^9^ CFU/mL, and boiled for 5 minutes. To isolate secreted proteins, Halt Protease Inhibitor (ThermoFisher) was added to *S. flexneri* supernatant which was then concentrated 15-fold using a 10 kDa molecular weight cutoff filter (Amicon). Concentrated supernatant was normalized to the OD_650_ of the bacterial culture from which they were collected and then resuspended in 4X Laemmli SDS sample buffer.

Samples were run on a 10% SDS-PAGE gel and then either stained with Coomassie Brilliant Blue or transferred to a 0.45 µm-pore-size nitrocellulose membrane (GE Healthcare) for immunoblotting with mouse monoclonal anti-S-peptide antibody (Thermo Fisher) diluted 1:1,000, mouse monoclonal anti-6x-His (Thermo Fisher) diluted 1:1,000, rabbit polyclonal anti-SecA antibody (Donald Oliver, Wesleyan University) diluted 1:10,000, mouse monoclonal anti-IpaC (Edwin Oaks, Walter Reed Army Institute of Research) diluted 1:300, or monkey anti-*S. flexneri* convalescent-phase antiserum (Edwin Oaks, Walter Reed Army Institute of Research) diluted 1:1,000. Blots were developed with a Pierce ECL detection kit (Thermo Fisher).

RNA isolation: *S. flexneri* was grown in the indicated growth medium to late log phase. Four mL of late log phase cell suspension was mixed with 1 mL of an ice-cold solution containing 95% ethanol and 5% phenol (pH 4.5), and then kept on ice until ready for further processing. Once all samples were ready, the cells were pelleted by centrifugation, resuspended in 100 µL of 1 mg/mL lysozyme in TE buffer, and incubated at room temperature for 5 minutes. 1 mL of RNA-Bee (Tel-Test Inc.) was added to the lysozyme-treated cells, 200 µL of chloroform was added, and then the solution was centrifuged at 4°C, 21,400 x *g* for 15 minutes. The aqueous phase was collected, mixed with an equal volume of isopropanol, and stored at -80°C overnight. Samples were centrifuged at 21,400 x *g* for 20 minutes to pellet precipitated RNA, which was then washed once with ice-cold 75% ethanol, air dried, resuspended in water, and DNase treated as per the manufacturer’s instructions (Turbo DNA-*free*, Invitrogen).

cDNA synthesis and quantitative PCR (qPCR): 2 µg of RNA was reverse transcribed into cDNA using the Superscript III kit (Thermo Fisher) as per the manufacturer’s instructions. Primers for qPCR were designed using Primer3 (http://bioinfo.ut.ee/primer3-0.4.0/). For Real-time qPCR, cDNA was diluted 1:10 and used with Power SYBR Green (Thermo Fisher). The qPCR was run on an Applied Biosystems ViiA7 instrument as previously described (81). Relative expression of virulence genes was calculated by using the ΔΔCt method and normalized to the mean of two reference genes; *secA* and *gyrA*.

Fractionation of *Bt* CM: *Bt* CM was fractionated by size using 10 kDa and 100 kDa molecular weight cutoff centrifugal filters (Amicon). Filters were rinsed once with sterile saline. Then 2.5 mL of *Bt* CM or BHIS was added to the filter and centrifuged at 3,000 x *g* at 4°C until all but 100 µL of liquid had flowed through the filter. The flowthrough was mixed 1:1 with BHIS, while the top fraction was resuspended in 5 mL of BHIS for *S. flexneri* growth. For ultracentrifugation, 2.5 mL of BHIS and *Bt* CM were centrifuged at 135,000 x *g* for 2 hours. The supernatant was collected. The resulting pellet was resuspended in 1 mL of sterile saline and centrifuged a second time. The supernatant was mixed 1:1 with fresh BHIS, while the pellet was resuspended in 5 mL of fresh BHIS for *S. flexneri* growth.

Proteinase K treatment of *Bt* CM: BHIS and *Bt* CM were treated with 50 µg/mL of Proteinase K at 37°C for 3.5 hours. Proteinase K treated BHIS and *Bt* CM were then each mixed 1:1 with fresh BHIS for *S. flexneri* growth.

Isolation and normalization of total lipids: Stationary phase *Bt* was pelleted by centrifugation for 10 minutes at 13,000 x *g*. Alternatively, large scale preps of crude *Bt* OMVs were isolated by centrifuging 75 mLs of *Bt* CM for 3.5 hours at 38,400 x *g*. Total *Bt* lipids from either cell pellets or crude OMVs were extracted by the method of Bligh and Dyer (82). For a negative control, the crude OMV isolation and lipid extraction was performed starting with 75 mLs of uninoculated BHIS to generate a sample referred to as “Mock lipids”. *E. coli* total lipid extracts were purchased (Avanti Polar Lipids). Extracted or purchased lipids were dried under a stream of nitrogen and then stored at -20°C until use.

For relative quantitation, lipids were resuspended in saline and then normalized by their fluorescence in the presence of the lipophilic fluorescent dye FM4-64 FX by a method adapted from a previously published protocol (83). Briefly, two-fold serial dilutions of the lipid samples were made, 50 µL of each dilution was mixed with 150 µL of 6.67 µg/mL FM4-64 FX (final concentration 5 µg/mL) and incubated for 10 minutes at room temperature. Fluorescence was measured at (Ex/Em, 510/640 nm) on a FlexStation 3 plate reader. Serial dilutions falling within the linear range of the assay were used to determine relative concentrations of the lipid preparations. To standardize lipid concentrations, lipid samples were adjusted with saline, such that a 50 µL aliquot gave a signal of 240 relative fluorescent units (RFU) in a 200 µL reaction (5 µg/mL FM4-64 FX). To assess the effect of lipids on *S. flexneri* IpaC levels, normalized lipid stocks (240 RFU) were diluted 6-fold in *S. flexneri* growth medium for a final lipid concentration of 40 RFU.

Outer Membrane Vesicle (OMV) Isolation: To isolate OMVs, 240 mL of *Bt* CM, prepared as described above, was concentrated to 7 mL using either 100 kDa cutoff centrifugal filters (Amicon) or a tangential flow filtration device with a 100 kDa cutoff filter (Vivaflow 50R, Sartorius). Concentrated *Bt* CM was then ultracentrifuged at 135,000 x *g* for 2 hours to pellet crude *Bt* OMVs. Depending on the downstream application, the crude OMVs were either washed once with saline or further purified on an OptiPrep (Sigma) density gradient modified from a published protocol (84). Briefly, the crude OMVs were resuspended in 45% OptiPrep [vol/vol] and added to the bottom of an Ultraclear (Beckman) centrifuge tube. Five layers of OptiPrep, decreasing in increments of 5% each layer down to 20% OptiPrep, were layered on top of the OMVs. The OMVs were then ultracentrifuged at 4°C for 20 hours at 150,000 x *g*. The density gradient was collected in 12 fractions and SDS-PAGE was performed to identify the fractions that contained OMVs. OMV-containing fractions were then pooled, diluted in 10-fold excess Dulbecco’s phosphate buffered saline supplemented with salts (DPBS) (0.2 g/L KCl, 0.2 g/L KH_2_PO_4_, 11.7 g/L NaCl, 1.15 g/L Na_2_PO_4_, 0.1 g/L MgCl_2_·6H_2_O, and 0.1 g/L CaCl_2_), and the OMVs were collected by centrifugation at 38,400 x *g* for 3.5 hours. The entire OMV isolation protocol was also performed on 240 mL of sterile BHIS to generate a sample for negative controls, referred to as mock extract.

Inner membrane/Outer membrane preps: To isolate inner and outer membranes (IM and OM, respectively), a stationary phase culture of the indicated *Bt* strain was centrifuged at 13,000 x *g* for 10 minutes. The resulting pellet was resuspended in buffer (10 mM Na_2_HPO_4_ and 5 mM MgSO_4_), lysed by sonication, and then centrifuged at 13,000 x *g* to remove cell debris. Total membrane (TM) was isolated by centrifuging the supernatant at 135,000 x *g* for 40 minutes. To separate inner and outer membranes, the total membrane pellet was resuspended in 1% (wt/vol) N-lauroyl sarcosine (sarkosyl) and incubated with rocking at room temperature for 1 hour. To separate IM from OM, the sample was centrifuged at 135,000 x *g* for 40 minutes. The sarkosyl-soluble supernatant, which contained the inner membrane fraction, was collected. The sarkosyl-insoluble pellet, which contained the outer membrane, was washed with 1% sarkosyl, centrifuged at 135,000 x *g* for 40 minutes, and then resuspended in water for downstream applications.

Quantification of Membrane Proteins: For quantification of protein, total membranes, outer membranes or OMVs were resuspended in 0.5% Triton X-100, while inner membranes were resuspended in 1% N-lauroyl sarcosine. Protein concentration was quantified by the DC Protein Assay (Bio-Rad) as per the manufacturer’s instructions. For all experiments involving OMVs, OMV concentrations are normalized by the total protein concentration of the vesicles.

Transmission Electron Microscopy: Five µL of OMVs that had been diluted 1:10 in saline were added to a glow discharged 200 mesh formvar/carbon grid (Electron Microscopy Sciences, FCF200-Cu) and incubated for 5 minutes. Excess sample was wicked off, the grid was washed once with a drop of water, and stained for 1 minute with 2% uranyl acetate before being air dried and imaged. Imaging was performed on a FEI Tecnai Spirit TEM at the Center for Biomedical Research Support Microscopy and Imaging Facility at UT Austin (RRID# SCR_021756)

Expression of His-tagged proteins in *Bt*: *Bacteroides* expression vector pFD340 (85) expressing His-tagged membrane proteins was transformed into the donor strain *E. coli* S17-1 *λpir* and conjugated into *Bt* as previously described (86). Briefly, pellets from overnight cultures of *E. coli* S17-1 *λpir* expressing the tagged gene of interest and *Bt* were pooled, plated on BHI agar plates, and grown aerobically overnight at 37°C without selection. The resulting lawn was resuspended in a small volume of BHIS. To recover single colonies of *Bt* conjugants, dilutions were plated on BHI agar containing gentamicin and erythromycin and incubated anaerobically at 37°C for 24 hours.

OMV uptake assay: To assess fusion of *Bt* OMVs by *S. flexneri*, a slight modification of a previously published protocol was used (87). Briefly, 50 µg of purified *Bt* OMVs in 250 µL of saline was stained with 5 µg/mL of FM4-64 FX at room temperature for 15 minutes. To eliminate excess unbound dye, the stained OMVs were washed three times with a 20-fold excess volume of saline on a centrifugal filter (100 kDa molecular weight cutoff, Amicon). To control for dye carryover, the staining and washing procedure described above was also performed on 250 µL of saline in the absence of OMVs. To measure OMV uptake, *S. flexneri* from a late log phase culture was adjusted to an OD_650_ of 1.0 in 1 mL of saline and co-incubated, rocking at room temperature, with either 50 µg of stained OMVs or an equal volume of stained saline in the absence of OMVs. At the indicated time points, 500 µL of sample was removed, washed three times with saline, fixed for 10 minutes in 4% paraformaldehyde, and washed three times more. The fluorescence (Ex/Em of 510/640 nm, SpectraMax M3) and OD_650_ (FlexStation 3) of *S. flexneri* that had been co-incubated with stained OMVs was read on the respective microplate readers. To control for dye diffusion from OMVs, the OMVs were stained as described above and then soaked in 1mL of saline for 30 minutes. After 30 minutes the soaked OMVS were separated by centrifugation at 135,000 x *g* for 1 hour. The supernatant and soaked OMVs were each collected and used in an OMV uptake assay as described above.

## Acknowledgements

This work was supported by grant AI016935 from NIAID. We thank Mario Feldman for the generous gift of plasmids pFD340::BT_0148 and pFD340::BT_1488 and Donald Oliver for the anti-SecA antibody. We thank Alexandra Mey and Carolyn Fisher for thoughtful discussion and critical reading of the manuscript, and Bryan Davies for assistance with anaerobic cell culture. We thank Michelle Mikesh, Center for Biomedical Research Support Microscopy and Imaging Facility at UT Austin (RRID# SCR_021756) for assistance with TEM. Figures 3A and 4A were created with BioRender.com.

## Supplement legends

Table S1. Strains and plasmids used in this study.

Table S2. Primers used in this study.

Figure S1: *Bt* CM reduces *S. flexneri* virulence factor secretion. Secreted proteins were collected from *S. flexneri* grown in either BHIS or ½ *Bt* CM. A Western blot using monkey anti-*Shigella* convalescent-phase antiserum was performed to look at the relative secretion of a panel of virulence factors. A representative Western blot of 5 independent replicates is shown.

Figure S2: *E*.*coli* CM does not affect *S. flexneri* virulence gene expression. Relative virulence gene expression of *S. flexneri* grown to log phase in either BHIS or ½ *E*.*coli* CM was measured by RT-qPCR. The *E. coli* CM was tested in parallel with the BT CM shown in Fig. 3B, and the BHIS control values are the same as shown in that figure. *C*_*t*_ values were normalized to the mean of two endogenous controls, *gyrA* and *secA. P* values were determined from the Δ*C*_*t*_ values of three biological replicates using a two-tailed Student’s *t* test with Holm-Šídák correction for multiple comparisons. Error bars indicate SD.

Figure S3: *Bt* OMVs have the expected protein profile: BT_0418 is localized to the outer membrane, while BT_1488 is localized to outer membrane vesicles. Total membrane (TM), inner membrane (IM), outer membrane (OM) or outer membrane vesicles (OMV) were extracted from *Bt::*pFD340/BT_0418-6xHis and *Bt*::pFD340/BT_1488-6xHis strains. Membrane preps were normalized using a DC protein assay, and 10 µg of sample was loaded per well for an anti-His Western blot.

Figure S4: Direct contact between *Bt* OMVs and *S. flexneri* is required for dye transfer. *Bt* OMVs were stained with 5 µg/mL of the fluorescent dye FM4-64 FX, washed extensively, soaked in saline for 30 minutes, and the separated out of their soaking solution. *S. flexneri* was incubated in the OMV soaking solution, the presence of soaked OMVs, or the presence of unsoaked stained OMVs. At a timepoint 2 hours, *S. flexneri* fluorescence was measured on a plate reader (Ex/Em of 510/640 nm). *P* values were determined from three biological replicates using a one-way Anova with Dunnett’s test for multiple comparisons (*, *p* < 0.05). Error bars indicate standard error.

